# Avian elevational migrants in India’s Western Ghats challenge climatic constraint theory through broader environmental tolerances

**DOI:** 10.1101/2024.12.14.628488

**Authors:** V.A. Akshay, Caitlin J. Campbell, Bette Loiselle, Robert Guralnick

## Abstract

1. Avian elevational migration is a common but often overlooked form of short-distance migration. Numerous hypotheses have been proposed to explain this behavior, yet few studies have tested those comprehensively. We examine the climatic constraint hypothesis – a key hypothesis explaining elevational migration – to evaluate its predictions for avian elevational migration in the monsoon-dominated Western Ghats of India.
2. We used citizen science data from eBird to quantify elevational migration patterns of resident birds of the region. Using robust data curation protocols and new bias correction methods, we modeled migration patterns of 164 species across summer, monsoon and winter seasons. To formally test the climatic constraint hypothesis, we modeled the likelihood and distance of migration against body size and environmental niche breadths, using phylogenetically informed logistic regression and Bayesian generalized linear mixed models.
3. Our results show that 43% of resident birds exhibited elevational migration in at least one season. Most species migrated between winter and summer (57), followed by migrations between summer and monsoon (39) and then monsoon and winter (38). Species predominantly moved upslope in summer (82%) and downslope in monsoon (87%), with no discernible pattern of direction in winter. Species with broader temperature tolerances were more likely to migrate across all seasons. Additionally, species with broader precipitation niches and larger body sizes moved further downslope during the monsoon and winter, respectively.
4. We found that species with broader tolerances to environmental conditions were more likely to be migratory and moved further along the elevational gradient, contradictory to the expectations of the climatic constraint hypothesis. Our analyses suggest that elevational migrants of the Western Ghats are not climatically limited but likely possess flexibility to track ecological resources that vary with season and elevation.

## Introduction

Animal migration is one of nature’s most spectacular phenomena, yet we are just beginning to understand the patterns of migration and underlying eco-evolutionary dynamics that determine those patterns. Migration is thought to be favored when the fitness benefits of exploiting spatiotemporally varying resources outweigh the associated costs (Zurell et al., 2018). Migrating animals can take advantage of multiple habitats where suitability (e.g., survival, food availability, predation risk, and reproductive success) differs across seasons and regions (Hsiung et al., 2018; Milner-Gulland et al., 2011). Migration also involves significant costs, such as increased energy expenditure, heightened predation risk during transit, and potentially shorter breeding seasons due to time spent migrating (Alves et al., 2013; Boyle, 2017; Milner-Gulland et al., 2011). These trade-offs influence the evolution of migratory behaviors, where organisms optimize their fitness by balancing the rewards of resource acquisition with the risks and energy costs associated with movement (Bowlin et al., 2010; Shaw, 2016).

While some migrants travel very long distances, many make shorter journeys. Short-distance migrants commonly perform seasonal back-and-forth movements between regions with altering favorable and unfavorable conditions, including breeding areas. Short-distance migrants are often less time-constrained, thus able to focus on minimizing energetic costs during their seasonal transitions (Dingle & Drake, 2007; Rappole, 2013). Elevational migration is a form of short-distance migration where animals move between breeding and non-breeding grounds that differ in elevation in a cyclical annual pattern (Rappole, 2013). In birds, elevational migration is relatively common, with ∼10% of species showing seasonal elevational shifts (Barçante et al., 2017). Although a majority (44%) of avian elevational migrants occur in the neotropics (Barçante et al., 2017), elevational migration occurs worldwide and is documented in ∼ 20% of continental North American birds (Boyle, 2017), 55–65 % of breeding Himalayan species (Menon et al., 2023), and ∼58% of resident Taiwanese species (Tsai et al., 2021).

The underlying intrinsic and extrinsic mechanisms explaining elevational migration in birds has been a focus of ornithological research, especially recently (Williamson & Witt, 2021). This has resulted in numerous non-exclusive hypotheses to explain these seasonal elevational shifts. We note three key ones that have garnered empirical support. The first is the ‘climatic constraint hypothesis;’ where seasonal climatic regimes can expose individuals to unsuitable temperatures, precipitation, and wind speeds at their breeding elevations during the non-breeding season, thus, imposing strong constraints on a species’ ability to maintain optimal physiological conditions (Hsiung et al., 2018). To minimize these direct weather-driven physiological constraints, species can move to elevations with more suitable conditions. The central expectation of this hypothesis is that species with narrower tolerances to harsh weather conditions (i.e. those with narrower climatic niche breadths or smaller body size) will move further downslope during the winter season and upslope during the breeding season to track suitable climate conditions. Recent work by Tsai et al. (2021) and Menon et al. (2023) support this hypothesis and have demonstrated that Taiwanese and Himalayan birds that are more cold intolerant make larger elevational migrations between their breeding and non-breeding grounds.

Another set of hypotheses, related to food resources, stem from formative work on avian elevational migrants in the neotropics. The ‘food availability hypothesis’ posits that bird species move along elevational gradients to track seasonally varying food resources (Levey, 1988; Loiselle & Blake, 1991; Stiles, 1988). Studies on frugivorous birds in Costa Rica showed a strong correlation between peak fruiting season and elevational distribution of the study species, adding support to this hypothesis (Levey, 1988; Loiselle & Blake, 1991). Later tests of the ‘food availability hypothesis’ in Costa Rica (Boyle, 2008b, 2011; Boyle et al., 2010) suggested it is likely limited foraging opportunities rather than food resources, per se, that lead to elevational migrations. In particular, the ‘limited foraging hypotheses’ (LFO) posits that heavy wet-season storms cause weather-driven reductions in foraging opportunities, driving taxa with higher foraging needs to migrate to avoid starvation (Boyle, 2008b). Initial tests of the LFO hypothesis on the White-ruffed Manakin (*Corapipo altera*) in Costa Rica showed increased captures of elevational migrants at lower elevations following heavy rainfall (Boyle et al., 2010). Community-level studies later confirmed that storms at higher elevations led to a higher abundance of migrants in the lowlands, with body size and diet influencing responses (Boyle, 2011). These studies suggest that species will shift locations to maximize foraging success during periods of poor conditions, indicating an adaptive flexibility in response to changing conditions. While empirical support exists for the hypotheses outlined above, testing of their expectations on avian elevational migrants remains limited to shifts between breeding and non-breeding grounds, with little attention to movements over complete annual cycles (but see Rueda-Uribe et al., 2024). Elevational movements may be more complex, particularly in highly seasonal, tropical montane systems where challenging monsoonal conditions can lead to unique survival challenges. Tropical birds may therefore make multiple movements across a year in response to time-varying environmental conditions and associated changes in ecological resources and habitat quality. Improving our understanding of elevational movements of birds across complete annual cycles could lend further support to traditional hypotheses explaining this complex and understudied behavior.

A significant challenge in studying avian elevational migration over annual cycles is the need for year-round monitoring over broad geographic extents. Citizen science initiatives, such as eBird, can help overcome some of these challenges via assembling semi-structured checklists in a common format submitted by an extensive network of citizen scientists (Kelling et al., 2019; Sullivan et al., 2009). Still, these data resources, however extensively verified or complete, contain sources of bias including uneven sampling over space and time (Sullivan et al., 2014). In the context of avian elevational migration, biases in citizen science data can skew results towards elevations where sampling has been more frequent. Researchers have employed various statistical methods to correct these biases, specifically resampling bird occurrences within elevational bands to obtain more uniform sampling across elevations (Menon et al., 2023; Tsai et al., 2021). Yet, there is scope for further improvement in correcting biases from semi-structured checklists, particularly to shift from case-specific methods to a more generalizable framework.

In this study we used bird occurrence data obtained from the eBird portal to quantify elevational migration of resident birds in the Western Ghats of India and explicitly test the expectations of the climatic constraint hypotheses as an explanatory framework of this behavior. Support or falsification of this key hypothesis is a first step towards addressing broader questions of determinants of seasonal migration in highly monsoonal tropical systems. Our efforts are focused on the Western Ghats, which are a 2200 km long mountain range extending along the western coast of the Indian subcontinent, covering tropical latitudes of 8 N - 21 N (Das et al., 2006); and Figure 1a) and form a sizable portion of the Western Ghats and Sri Lanka biodiversity hotspot (Myers et al., 2000). These mountains are topographically, climatically, and biologically diverse (Gadgil & Meher-Homji, 1990, Pascal, 1988) and include an elevational gradient of 40 - 2625 m (Figure 1a), with significant temperature variation between the plains and higher elevations (Figure 1b). This mountain range also experiences extreme monsoon conditions during June to October due to the annual south-west monsoon (Figure 1d & 1e). Broadly, the Western Ghats experience three distinct seasons: a hot and dry “summer” from March to May, a cool and extremely wet monsoon from June to October and a cool and dry “winter” from November to February (Figure 1c & 1e). These seasonally variable conditions may pose significant challenges to resident birds, necessitating year-round elevational movements to adapt to changing seasonal dynamics.

**Figure 1:**
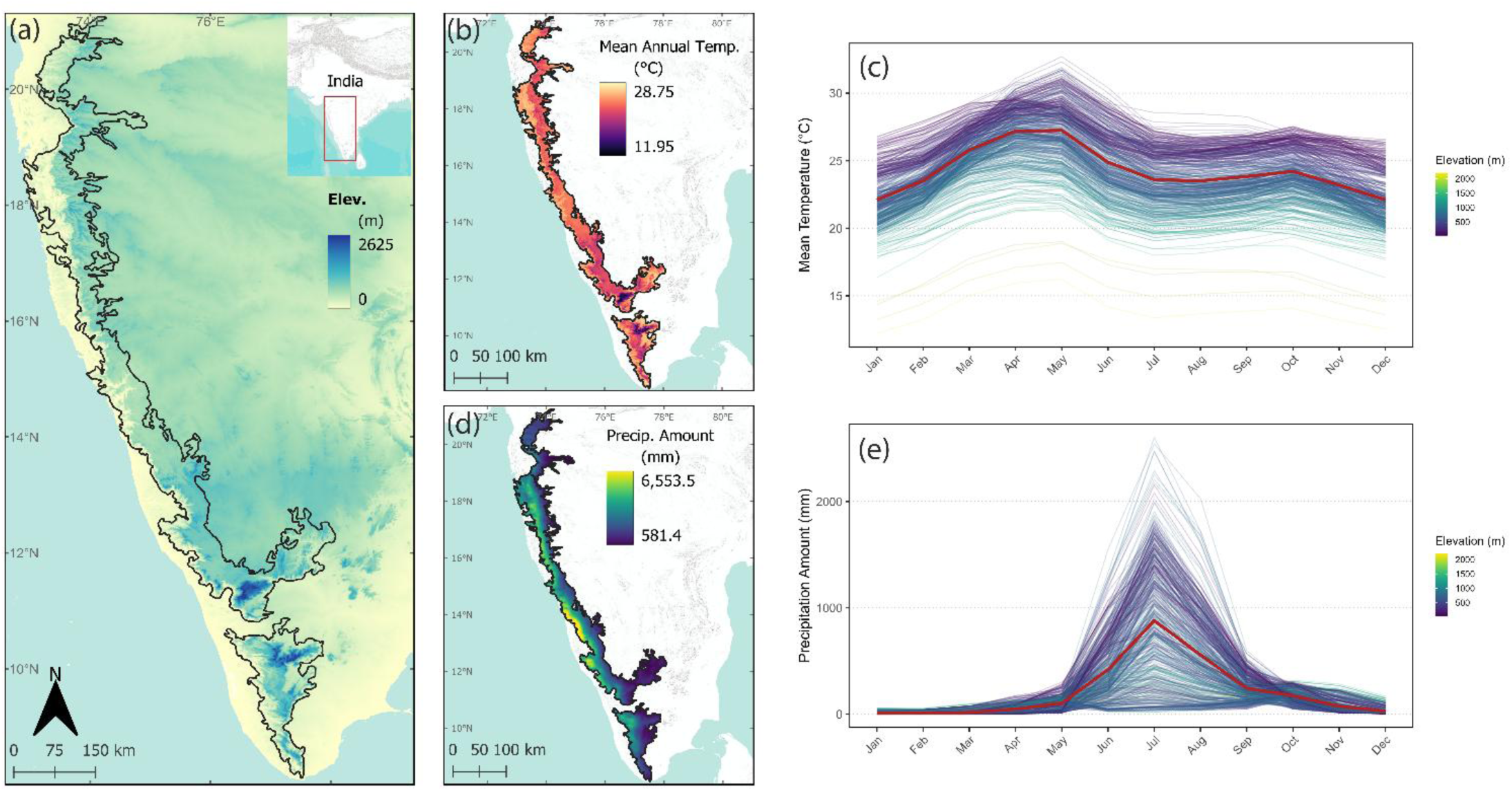
The Western Ghats of India (boundary shown as black line) are topographically and climatically unique. (a) Latitudinal extent and elevational range of the Western Ghats. (b) Mean annual temperature in the region ranges from 12°C at higher elevations to 25°C in the plains. (c) Temperatures are highest during the summer season from March to May across the elevational gradient (red line indicates mean temperature across the year). (d) The Western Ghats creates a strong rain shadow effect, leading to a longitudinal precipitation gradient. (e) Precipitation peaks during the months of June to October during the annual south-west monsoon (red line indicates mean precipitation across the year). Elevation is built from 30 seconds resolution SRTM data (Farr et al., 2007) and climatic variables are visualized from CHELSA data at 1 km resolution (Karger et al., 2017).

Here, our overall goals are to (1) quantify elevational migration distance and identify patterns of movement of resident birds of the Western Ghats over a complete annual seasonal cycle (summer, monsoon and winter), and (2) test the effect of a species’ body size, and environmental niche breadths on their likelihood of being a seasonal elevational migrant and elevational migration distance. We expect species to demonstrate upslope movements during the summer breeding season and downslope movements in the monsoon and winter seasons as has been demonstrated in other tropical elevational migrants. Further we expect that small-bodied species with narrower environmental niche breadths will be more likely to be elevational migrants and will make longer distance seasonal elevational movements, consistent with the climatic constraint hypothesis. Finally, we present a generalizable method for correcting sampling biases in citizen science data and provide evidence of its effectiveness compared to previous methods.

## Materials and Methods

### eBird occurrence records and data filtering

We downloaded eBird checklist data from India during the time period from 2013 - 2023. We first applied a spatial filter to these data to include checklists that fell within the bounds of the Western Ghats. Next, we filtered these data to include records between March 2013 and March 2023, yielding 10 full years of occurrence data. The ten years of data were aggregated for all further analysis. We retained occurrence records associated with complete checklists, where observer(s) recorded all identifiable species. To standardize sampling protocol, we included checklists that followed the “stationary” or “traveling” protocols based on the distance moved by the observer. To minimize extreme variation in sampling effort, checklists with a sampling duration of >5 hours, sampling distance of >1 km and with >10 observers were excluded (following Strimas-Mackey et al., 2020 and Tsai et al., 2021). The dataset was further filtered to exclude long-distance migrants and limited to species belonging to the orders: *Passeriformes*, *Bucerotiformes*, *Trogoniformes*, *Piciformes*, *Coraciiformes*, *Cuculiformes*, *Psittaciformes*, and *Columbiformes*. We excluded raptors (*Accipitriformes* and *Falconidae*), swifts (*Apodiformes*) and swallows (*Hirundinidae*) since these groups are generally observed in-flight and associating elevation to these records is difficult (Menon et al., 2023; Ramesh et al., 2022). Checklists were assigned a value for season based on the observation month (summer: March – May; monsoon: June – October; winter: November – February) and elevation of each checklist was obtained from the ASTER Global Digital Elevation Model at 30 m resolution (Abrams et al., 2020). To account for geographic location uncertainty, elevations were averaged using a 100 m buffer around each checklist location (Tsai et al., 2021) and rounded to two decimal places. All data filtering was performed in the programming environment R v. 4.3.3 (R Core Team, 2024), using the packages ‘auk’ (Strimas-Mackey et al., 2023) and ‘sf’ (Pebesma, 2018).

### Bias correction and method validation

A key challenge of eBird data is the presence of spatial sampling biases (Sullivan et al., 2014). eBird checklists are often located closer to roads (Ramesh et al., 2022), lower elevations are sampled more often than higher elevations (Menon et al., 2023; Tsai et al., 2021) and sampling is frequently seasonally biased. We examined the distribution of occurrence records and checklists as a function of elevation and season to understand sampling effort across our dataset. Sampling was uneven across elevations and seasons: most occurrence records were reported from elevations below 1200 m, and sampling was significantly higher in the winter when conditions are most pleasant (Figure S1.1). To account for this bias, we developed a resampling approach that helps correct these issues. We first calculated the cumulative minutes of sampling for each season/elevation combination in our filtered dataset as a measure of sampling effort. We then fit Generalized Additive Models (GAMs; Hastie & Tibshirani, 1986) for each season to predict sampling effort across the elevational range of occurrence records in our dataset. GAMs were built using the ‘mgcv’ (Wood, 2017) package in the R programming environment. Next, we resampled each species – season combination 20,000 times, with replacement, weighted by the inverse of the predicted value of sampling effort. We used the inverse of sampling effort to up-sample elevations with lower effort and down-sample elevations with higher effort. This resampling approach provides continuous elevational distributions of each species per season under more equal sampling effort. Using this bias adjusted distribution we extracted each species’ median (50^th^ percentile) elevation per season. Species with fewer than 10 occurrence records for any one season were excluded.

To test the effectiveness of our approach, we validated our method against the effort correction protocol used in Tsai et al. (2021), who used a different resampling methodology. Briefly, those authors binned sampling effort across elevational band and season and resampled within those bins. In contrast, our approach smooths noise in sampling via fitted GAMs across a continuous range of elevations. Further, we took care to reduce the effect of outliers in our resampling approach by only including species with at least 10 occurrence records in each season. We replicated the analyses of Tsai et al. (2021) and then applied our method for bias correction on their data to obtain estimates of median elevation for each species in their study. In comparison, we found that our method produced consistent and reasonably conservative estimates of median elevation (Figure S1.2 & S1.3) and has distinct advantages with respect to execution time (Figure S1.4).

### Quantifying elevational migration

We used our method for bias correction on the filtered bird occurrence data and extracted median elevation values for all species in each season. We classified a species as an elevational migrant in a given season if the 95% confidence intervals (CIs) around the median elevation estimate did not overlap between that season and the previous one. This resulted in a binary variable for each season denoting whether a species was an elevational migrant. We then calculated elevational migration distance for a species as the difference between median elevation values between the focal and the previous season. This was repeated for all species and consecutive seasons to arrive at three elevational migration distances per species (Table S1.1). Notably, species could exhibit both statistically ‘significant’ and ‘non-significant’ elevational movements across seasons. A non-significant movement indicates that the elevation change falls within the CI of the previous season’s median, rather than implying no movement. For example, *Acridotheres fuscus* showed a significant upslope shift of 133 meters from winter to summer, followed by a significant downslope shift of 119 meters in the monsoon, and a non-significant 14-meter movement in winter (Table S1.1). According to our classification, this species would be identified as a migrant in summer and monsoon, but not in winter. Despite this classification, the cumulative change in elevation across the three seasons was close to zero (133 – 119 – 14 = 0), highlighting that elevational migration, as defined in this study, represents a round-trip pattern when assessed across the annual cycle.

### Species trait data

We calculated each species’ temperature, precipitation, and wind speed niche breadth within our filtered Western Ghats dataset. Having excluded long-distance migrants from our dataset, we considered each species’ population to be closed within the bounds of the Western Ghats. Thus, we estimated a species’ realized environmental niche breadth within the study systems as a proxy for their tolerance to environmental conditions. We understand that this approach may underestimate a species’ environmental niche breadth, however we were interested in their regional adaptations to climate and its associated effect on elevational movements. We used the mean monthly temperature, precipitation, and wind speed layers, downloaded from CHELSA (Karger et al., 2017), and masked them to the bounds of the Western Ghats. Due to a slight mismatch between the time period of occurrence records in our dataset (2013 - 2023) and CHELSA data availability (up to 2018), we used CHELSA layers from 2008–2018. Next, for each occurrence point of a species in our dataset, we extracted temperature, precipitation, and wind speed values across all ten years of the masked environmental layers. We did not match occurrence records to our monthly environmental layers, but rather time averaged temperature, precipitation and wind speed values across the temporal range of the environmental data. We then calculated each species’ 2.5^th^ and 97.5^th^ percentile values for temperature, precipitation and wind speed using the time averaged occurrence specific environmental data. The difference between the 97.5^th^ and 2.5^th^ percentile was considered as the species’ environmental niche breadth. We believe this approach provided us with a reasonable estimate of a species’ environmental niche breadth given the modest timing mismatch in occurrence records and environmental data. Additionally, we extracted body mass (as a proxy for body size) from the AVONET global dataset (Tobias et al., 2022) for use in subsequent analyses.

### Statistical analyses

To identify the patterns of movement of species in our study, we used a chi-square test to evaluate the intra-seasonal variation in movement direction (upslope vs downslope). For each season, we calculated the observed frequencies of direction for species demonstrating significant elevational movement and compared these with expected frequencies of equal distribution between direction classes. Chi-square tests were performed using the ‘chisq.test’ function in R.

We then fit phylogenetic logistic regression models (Ives & Garland, 2010), separately for each season to understand the effect of body mass and environmental niche breadths on a species’ elevational migration status. We modeled elevational migration status (migrant or non-migrant) for a given season using a binomial distribution (see the “Quantifying elevational migration” section above). We employed a backward stepwise model selection approach to select the most parsimonious combination of predictor variables that explain the elevational migration status of a species in a season. We used a consensus (Lapointe & Cucumel, 1997) Western Ghats bird tree as the basis for determining phylogenetic autocorrelation in our logistic regression models (built using Jetz et al., 2012). The phylogenetic logistic regression model was built using the ‘phyloglm’ function from the ‘phylolm’ package (Ho & Ane, 2014) in R and stepwise model selection was performed using the ‘phyloglmstep’ function from the same package.

Finally, we fit Bayesian generalized linear mixed models to describe the effect of body mass and environmental niche breadths on elevational migration distance. These models were fit separately for upslope and downslope elevational migrants to understand differences in trait effects for these two migratory strategies. The response variable was migration distance modeled as a function of season (categorical, with summer as the reference level), body mass, and environmental niche breadths (temperature, precipitation, and wind), including interaction terms between season and each trait to assess seasonal variation in trait effects. To account for phylogenetic non-independence among species, we incorporated a phylogenetic random effect using a Gaussian process prior with a covariance matrix derived from our consensus Western Ghats bird tree. Models were fit using the ‘brms’ package (Bürkner, 2017, 2018) in R, employing Hamiltonian Monte Carlo sampling with 4 chains, each with 4000 iterations (2000 warmup), ensuring convergence (Rhat ≈ 1) and sufficient effective sample sizes. Model fit was evaluated via Bayesian conditional and marginal R² (Gelman et al., 2019). The reported posterior summaries include mean estimates and 95% credible intervals, parameters with credible intervals excluding zero were interpreted as meaningful effects.

## Results

### Patterns of elevational migration

We analyzed 946,282 occurrence records from 105,725 eBird checklists to quantify elevational migration of 164 resident bird species of the Western Ghats over three seasons. We found that 71 (43%) species showed significant elevational migration in at least one season. Summers had the highest number of elevational migrants with 57 (35%) species, of which 47 displayed upslope movement (Figure 2). This was followed by the monsoon with 39 (24%) species, of which 34 exhibited downslope movements (Figure 2). Winters had a marginally lower number of elevational migrants with 38 (23%) species and did not reveal a clear pattern of movement (Figure 2 & Figure S1.5). We identified only 12 (7%) species that exhibited significant movement in all three seasons; however 34 species were common between the summer and monsoon followed by 26 species between the winter and summer, and 15 species between the monsoon and winter.

**Figure 2:**
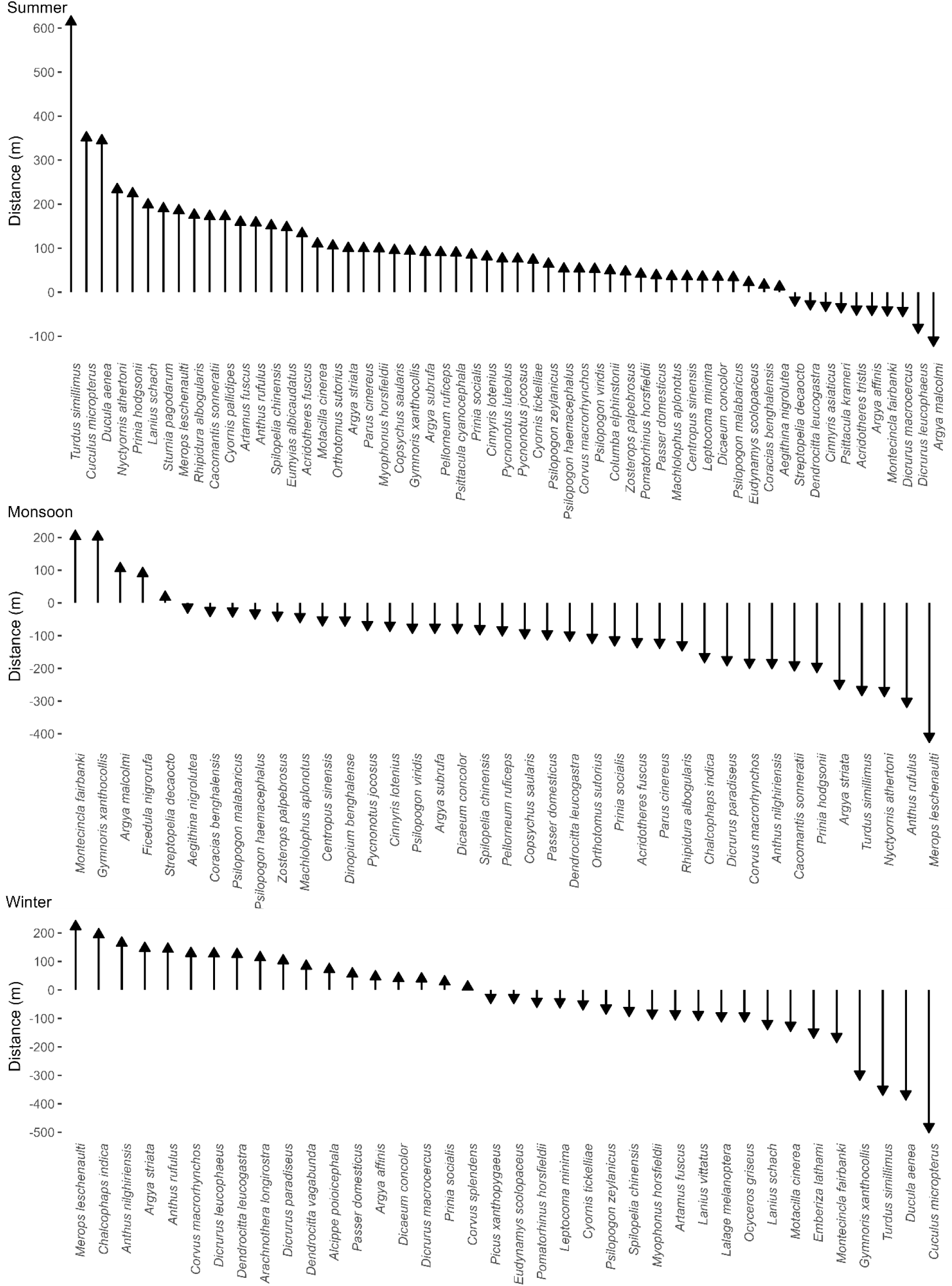
Patterns of elevational migration in this study for species that exhibit significant elevational movement between the seasons of interest. In the summer species moved predominantly upslope followed by downslope movement in the monsoon. The winter saw an almost even distribution of upslope and downslope elevational movements. The distance species move in meters (m) is represented as the length of the arrow.

More species demonstrated upslope movements in the summer and downslope movements in the monsoon than expected by chance (Chi-square tests; Table 1). There were no significant differences in movement direction during winter (Table 1).

**Table 1:**
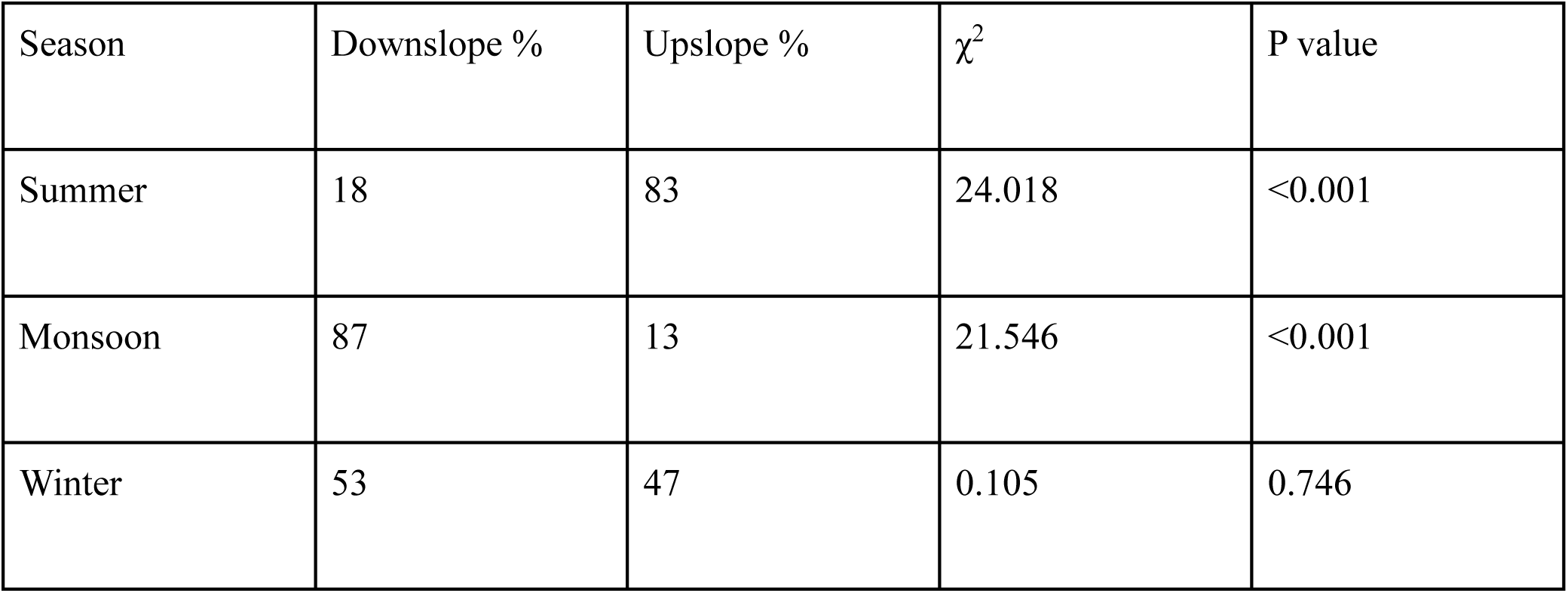
Chi-square test results examining the intra-seasonal variation in elevational migration direction of resident birds of the Western Ghats. Seasonal percentages of species demonstrating downslope (Downslope %) or upslope (Upslope %) movement are reported, along with chi-square statistics (χ^2^) and P values for each seasonal test.

### Association between species traits and elevational migration

Our backward model selection approach for the seasonal phylogenetic logistic regression models shows that temperature niche breadth was the strongest predictors of elevational migration status across all three seasons (Figure 3). Specifically, we found that elevational migration likelihood increased with broader temperature niche breadths regardless of season (Figure 3 & Table S1.2). Likelihood-based R² values indicated that the models provided a reasonable fit, explaining 14.7% of the variance in summer, 14.5% in monsoon, and 9.5% in winter beyond what was captured by the null model.

**Figure 3:**
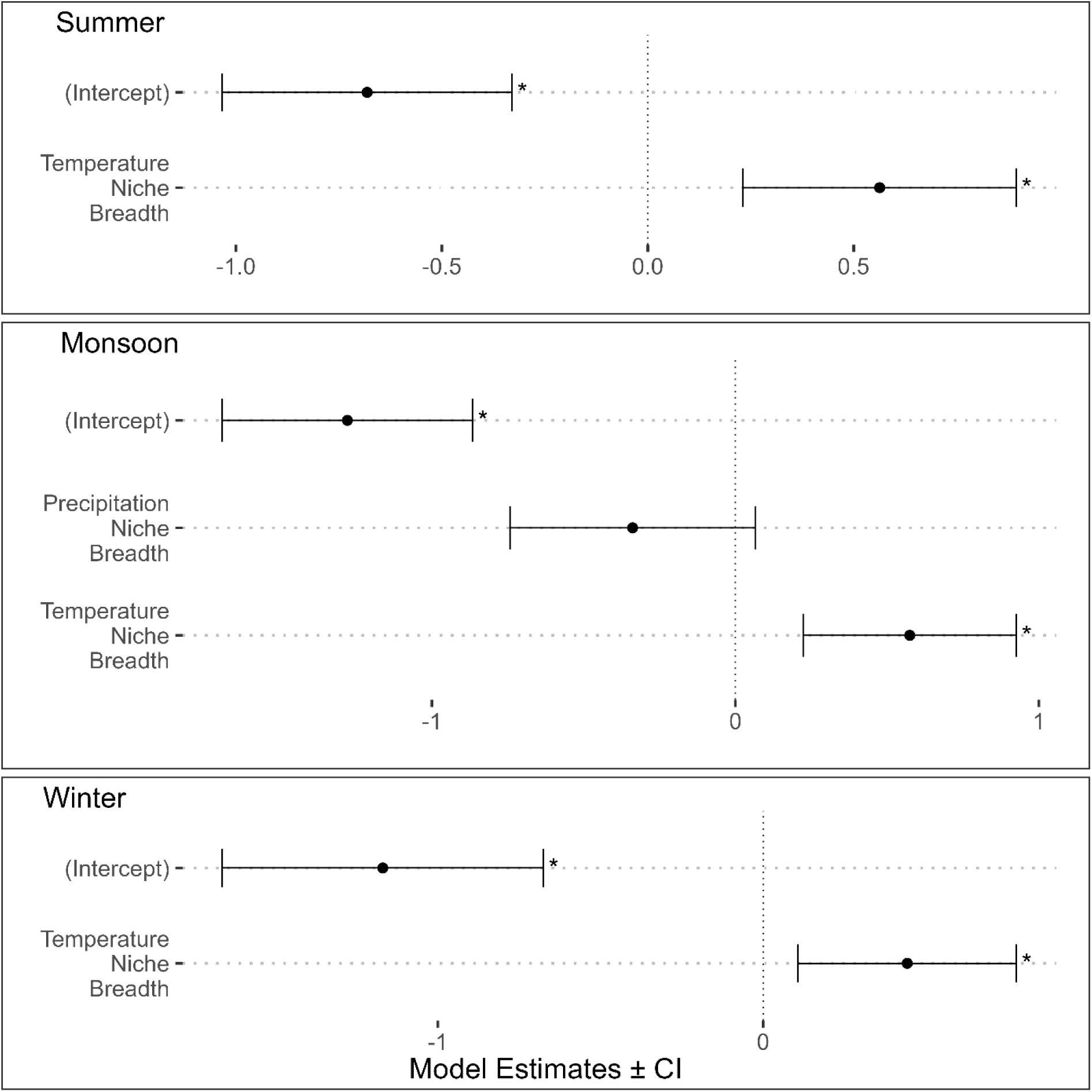
Parameter estimates for seasonal phylogenetic logistic regression models determining the effect of our candidate predictors on elevational migration status (migrant vs non-migrant). Dots represent model estimates with their 95% confidence intervals as error bars. Asterisk denotes significance (*p* < 0.05).

Elevational migration distance models, fit separately for upslope and downslope movement strategies, showed no significant trait effects for upslope movements in any season (Figure 4 & Table S1.3). In contrast, downslope models identified significant trait effects: species with broader precipitation niche breadths moved longer distances in the monsoon season and large-bodied species moved further downslope in the winter season (Figure 4 & Table S1.3). Model fit was reasonable, with the upslope model explaining 21.6% of the variance and the downslope model explaining 37.4%, including both fixed and random effects (Table S1.3).

**Figure 4:**
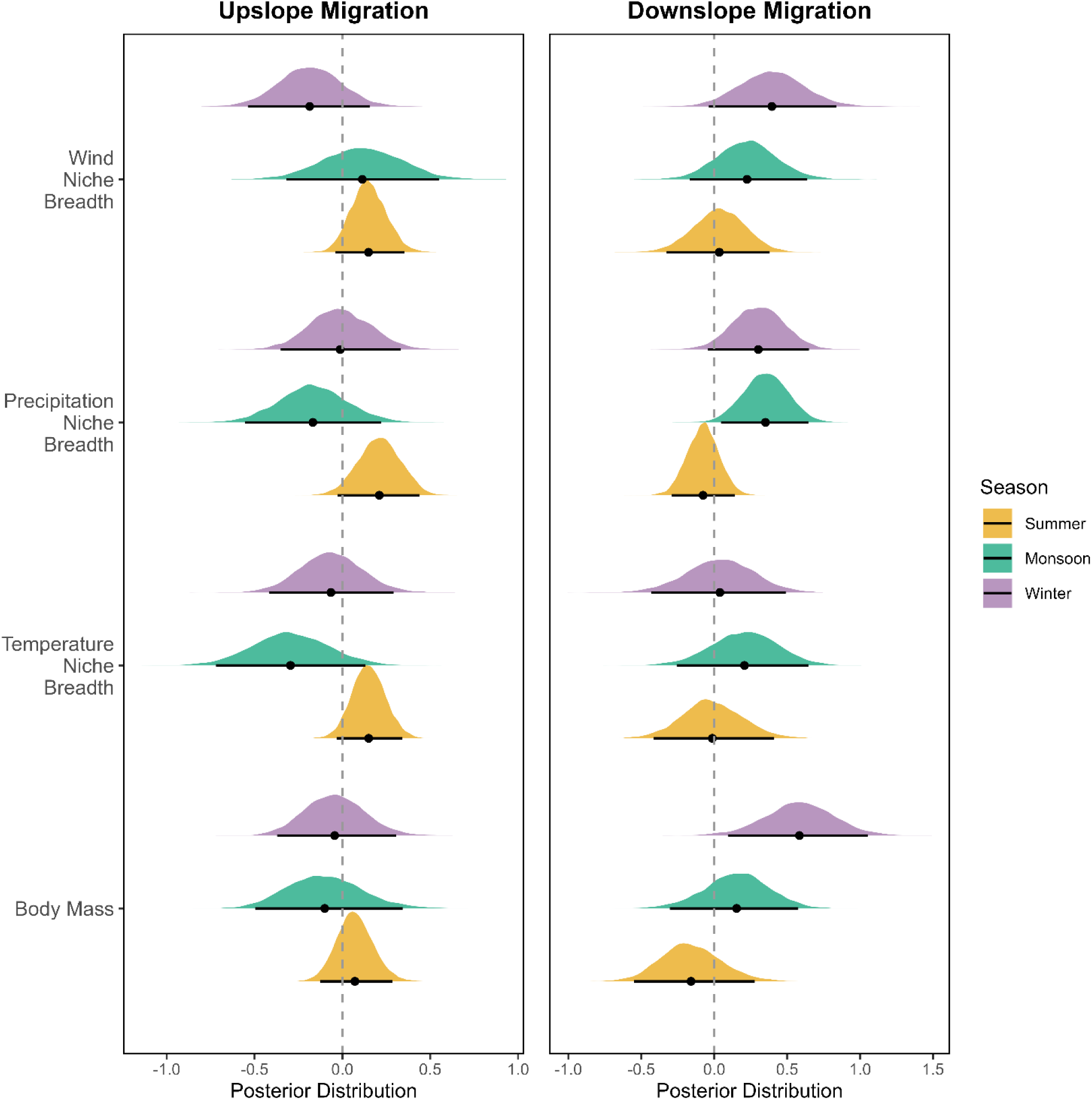
Posterior distributions of trait effects on upslope and downslope elevational migration distance across seasons in this study. Distributions represent estimates from Bayesian generalized linear mixed models testing the effect of body mass and environmental niche breadths on elevational migration distance. Posterior mean estimates are shown as dots with their 95% credible intervals as lines. Credible intervals that do not overlap zero indicate strong trait influence on migratory distance in a given season.

## Discussion

This study quantified the elevational migration strategies of 164 resident bird species in the Western Ghats of India. Utilizing a comprehensive citizen science dataset and robust analytical methods, we identified patterns of elevational movement and tested key predictions regarding this behavior. Our findings indicate that approximately 43% of resident birds exhibit seasonal elevational movements, a figure similar to other tropical regions (44% in Barçante et al., 2017). We found significant directionality in these movements, with species predominantly moving upslope during the summer breeding season and downslope during the monsoon, which is broadly consistent with predictions from the climatic constraint hypothesis but also applies to other hypotheses related to food resources and foraging as well. We next modeled the likelihood and distance of migration against predicted species traits, using phylogenetically informed logistic regression and Bayesian generalized linear mixed models. However, contrary to the key predictions of the climatic constraint hypothesis, we found that species with broader thermal niches were more likely to be elevational migrants, regardless of season. Moreover, species with broader precipitation niches and larger body sizes moved further downslope during the monsoon and winter, respectively. Across our analyses, a common theme emerged: species with broader environmental niches were more likely to migrate and do so over larger elevational distances. Thus, our trait-based analyses suggest that migratory species in the Western Ghats are not climatically limited per se but rather may be better adapted to exploit elevational variation in other seasonally fluctuating resources.

A key goal of this study was to determine if there were clear patterns in the direction of elevational movement in the Western Ghats, a system with three very different seasons. We identified a clear pattern of movement during the summer and monsoon seasons. In the breeding summer season, most elevational migrants moved upslope from their wintering elevations. Breeding upslope migrations are well documented in species worldwide (Boyle, 2017; Levey, 1988; Menon et al., 2023; Tsai et al., 2021). Furthermore, evidence suggests that nest predation rates may be lower at higher elevations, as observed in the neotropics (Boyle, 2008a; Skutch, 1985). Therefore, elevational migrants in the Western Ghats may be moving upslope during the breeding summer season to elevations where breeding success is likely to be higher.

In the monsoon season, we observed a marked downslope movement of birds from their breeding elevations, a response expected under both the climatic constraint and Limited Foraging Opportunities (LFO) hypotheses. Heavy and prolonged rainfall – a defining characteristic of the monsoon in the Western Ghats – likely imposes strong physiological constraints on species, driving them to move downslope in search of more favorable conditions. Additionally, intense rainfall events are known to reduce foraging opportunities for small-bodied consumers like birds and bats (Boyle, 2010; Boyle et al., 2020; Foster, 1974; Voigt et al., 2011). During the Western Ghats monsoon, intense rainfall likely limits foraging opportunities at higher elevations, prompting species to seek refuge at lower elevations where food resources may be more readily available, consistent with studies examining the elevational response of birds to heavy wet-season storms (Boyle, 2011, 2017; Boyle et al., 2010). Therefore, the effect of weather driven physiological constraints and reductions in foraging opportunities may be driving monsoon season downslope elevational movements of birds of the region.

This study failed to identify a pattern of elevational movement in the winter. Winter in the Western Ghats is the most temperate season, with favorable temperatures across the elevational gradient and limited precipitation. During this season, birds are likely responding to other drivers, like food resource phenology, that were not included in this study. In sum, our results suggest that both climatic factors along with seasonal constraints in foraging opportunity may both explain seasonal directionality of elevational migration in the Western Ghats.

The second goal of our study was to evaluate how species-level traits influence elevational migratory strategy – specifically, the likelihood of migrating and the distance of movement. While seasonal directionality may be shaped by immediate environmental cues, these strategies may reflect longer-term ecological and evolutionary filters acting on species traits. The climatic constraint hypothesis predicts that species with smaller body sizes and narrower environmental niche breadths will have a higher likelihood of being migratory and will make longer migrations. Our results contradict these expectations, we find that across all seasons, species with broader thermal niches were more likely to be elevational migrants. Additionally, we found a seasonal response among downslope elevational migrants: species with broader precipitation niches and larger body size moved further in the monsoon and winter seasons respectively. Collectively, these findings suggest that broader tolerances to environmental conditions may provide species with the flexibility to move in response to a wider range of ecological opportunities. In the highly seasonal and topographically complex landscapes of the Western Ghats, such flexibility could allow birds to track resources that shift in both space and time - including pulses of food availability, changes in vegetation structure, or the availability of favorable nesting conditions. Rather than being forced upslope or downslope by narrow climatic limits, species with broader thermal and precipitation niches may be better positioned to optimize finding the most favorable conditions that shift in elevation with the seasons. In this light, elevational migration, at least in this system, may reflect less of a response to climatic constraints, and more of an adaptive strategy that takes advantage of seasonal environmental and resource heterogeneity. Still, one limitation of our study is that we cannot directly examine foraging rates given this work is extremely intensive and challenging, especially over many species and across seasons in a complex environment. More direct testing of foraging is still needed in order to provide strong and direct, versus circumstantial, support of the LFO.

Taken together, our findings point to a more complex and dynamic pattern of elevational migration than is typically captured by classical hypotheses. Rather than discrete, bidirectional movements between fixed breeding and non-breeding sites, many species in the Western Ghats undertake seasonal elevational shifts that unfold over the course of the year. For instance, species such as the Indian Blackbird (*Turdus simillimus*) and Chestnut-headed Bee-eater (*Merops leschenaulti*) exhibited substantial elevational movements in all three seasons examined, while others like the Nilgiri Flycatcher (*Eumyias albicaudatus*) and Green Imperial-Pigeon (*Ducula aenea*) moved only in summer or in both summer and winter. These varied strategies, observed in a majority of species, suggest that elevational migration in climatically heterogeneous montane systems may be more continuous and seasonally responsive than a simple binary of breeding and non-breeding movement implies. These migrations may be shaped by ecological and environmental cues that vary across the year and are not fully captured by species-level traits. Indeed, our trait-based models accounted for only a modest portion of the variation in elevational migration likelihood and distance – with the best-performing models explaining just 37% of the variance. One plausible explanation for this unexplained variation is that elevational migration in the Western Ghats is influenced by fine scale phenological cues and shifting resource availability, mechanisms well-documented in long-distance latitudinal migrants (Both et al., 2010; Cohen et al., 2019; Mayor et al., 2017; Youngflesh et al., 2021). In this topographically and climatically complex region, pulses of fruiting, flowering, and arthropod emergence are likely tied to seasonal rainfall and temperature regimes and may move upslope or downslope as the year progresses. Birds may be responding to such phenological shifts – for example, the onset of monsoon rains triggering arthropod abundance, or fruiting events of key tree species – that dictate when and where food becomes available. Species with broader physiological or behavioral tolerances may be better equipped to track these ephemeral resource peaks, resulting in more flexible or extensive elevational movements. While such responses are often interpreted through the lens of climatic filtering, our findings suggest they may instead reflect more complex interactions between resource phenology, microclimate, and species-specific ecological flexibility.

### Caveats and Conclusions

This work relied solely on citizen science data collected from the Western Ghats by thousands of citizen scientists across India. Despite rigorous filtering, inherent biases persisted (Figure S1.1). We developed a bias-correction approach using fitted GAMs to predict sampling effort and resampling occurrence records based on these predictions. The strength of this method lies in resampling across a gradient of occurrence records, weighted by a user-controlled scale of sampling effort, resulting in a continuous distribution of occurrence records across the gradient of interest. While applied here to an elevational gradient, the method can be adapted to other gradients with prior estimates of sampling effort. We acknowledge that despite these correction methods, biases in citizen science data may still arise. For this reason, we did not attempt to estimate the elevational boundaries of a species in this study. Instead, we estimated median elevations of a species, which are relatively less prone to error. By using measures of central tendency, we believe our estimates of species’ seasonal median elevation were not influenced by latent sampling biases and were reliable enough for drawing inferences.

In conclusion, our study revealed the complex and seasonally dynamic nature of elevational migration in the Western Ghats. Birds in this region exhibit multiple shifts in elevation across the year, likely responding to fine-scale ecological and environmental cues that vary with season and elevation. These patterns move beyond conventional views of migration as a single annual event, pointing instead to a more flexible, year-round strategy shaped by tropical seasonality. Our findings highlight that species with broader environmental tolerances are better equipped to track ephemeral resources across seasons – a capacity that likely offers ecological resilience. The repeated seasonal use of elevational gradients by many species emphasizes the importance of conserving not only core breeding areas but also the full spectrum of elevational refuges. With rapid land-use change and invasive species affecting the Western Ghats (Arasumani et al., 2019; Lele et al., 2020), protecting habitat along this critical elevational gradient is essential for sustaining avian populations and preserving biodiversity in this globally significant hotspot.

## Acknowledgements

We would like to thank the citizen scientists and eBirders of India, without whom this work would not be possible. We would also like to thank Dr. Michael Belitz and Dr. Scott Robinson for methodological and theoretical discussion on this work. This work was partially funded by the NSF: RCN: Cross-Scale Processes Impacting Biodiversity (DEB-1745562) and the Biodiversity Institute at the University of Florida, for which we would like to thank Dr. Ana Carnaval and Dr. Pamela Soltis respectively.

## Author Contributions

V.A. Akshay, Robert Guralnick and Bette Loiselle conceived the ideas and designed the study methodology. V.A. Akshay curated and led the analysis of the data with significant contributions from Caitlin J. Campbell and Robert Guralnick. V.A. Akshay led the writing of the manuscript. All authors contributed to critically reviewing manuscript drafts and gave final approval for publication.

## Statement on Inclusion

Our study brings together scientists from the two countries, including a scientist whose home country is the country where the research was conducted. We have made a conscious effort to identify and cite relevant research from the region of study. We also acknowledge that the data used in this study was collected by a diverse group of citizen scientists in India, whose contributions were essential. As scientists based in the global North, we recognize the structural advantages and opportunities available to us, and we affirm that these dedicated citizen scientists merit greater recognition and support.

## Data Availability Statement

Raw data was downloaded from eBird and can be accessed by visiting: https://science.ebird.org/en/use-ebird-data/download-ebird-data-products. The source code and data sets utilized in this study are available on https://zenodo.org/records/17063921.

