## Supplementary Tables and Figures for "Avian elevational migrants in India’s Western Ghats challenge climatic constraint theory through broader environmental tolerances"

**Appendix 1**

Figure S1.1: Sampling effort in our filtered eBird dataset across three seasons (Summer, Monsoon, Winter). (a) density plot of the distribution of occurrence records by elevation in each season, solid black line denotes the median. (b) bar plot of the number of eBird checklists per season.

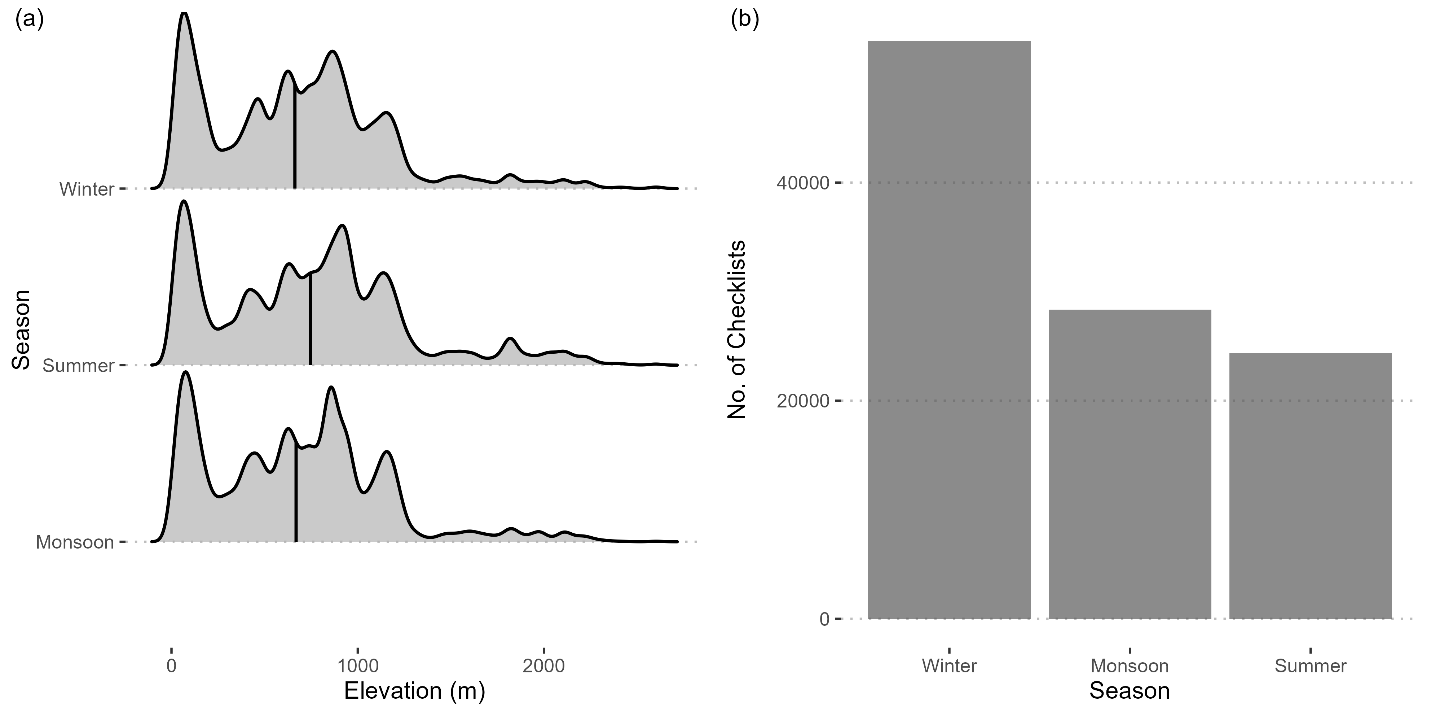

Figure S1.2: Scatter plot to compare the summer median elevation estimates of the resampling method employed in this study (Weighted Bootstrap) to the method employed by Tsai et al. (2021). Each panel is a comparison of our method to different levels of sampling effort and occurrence thresholds used by Tsai et al. (2021). Q1, Q2, Q3 denotes the first, second and third quantile of sampling effort, while 30 and 100 are the different occurrence thresholds in their analysis. A solid black 1:1 line is included to offer easier comparison between the two methods.

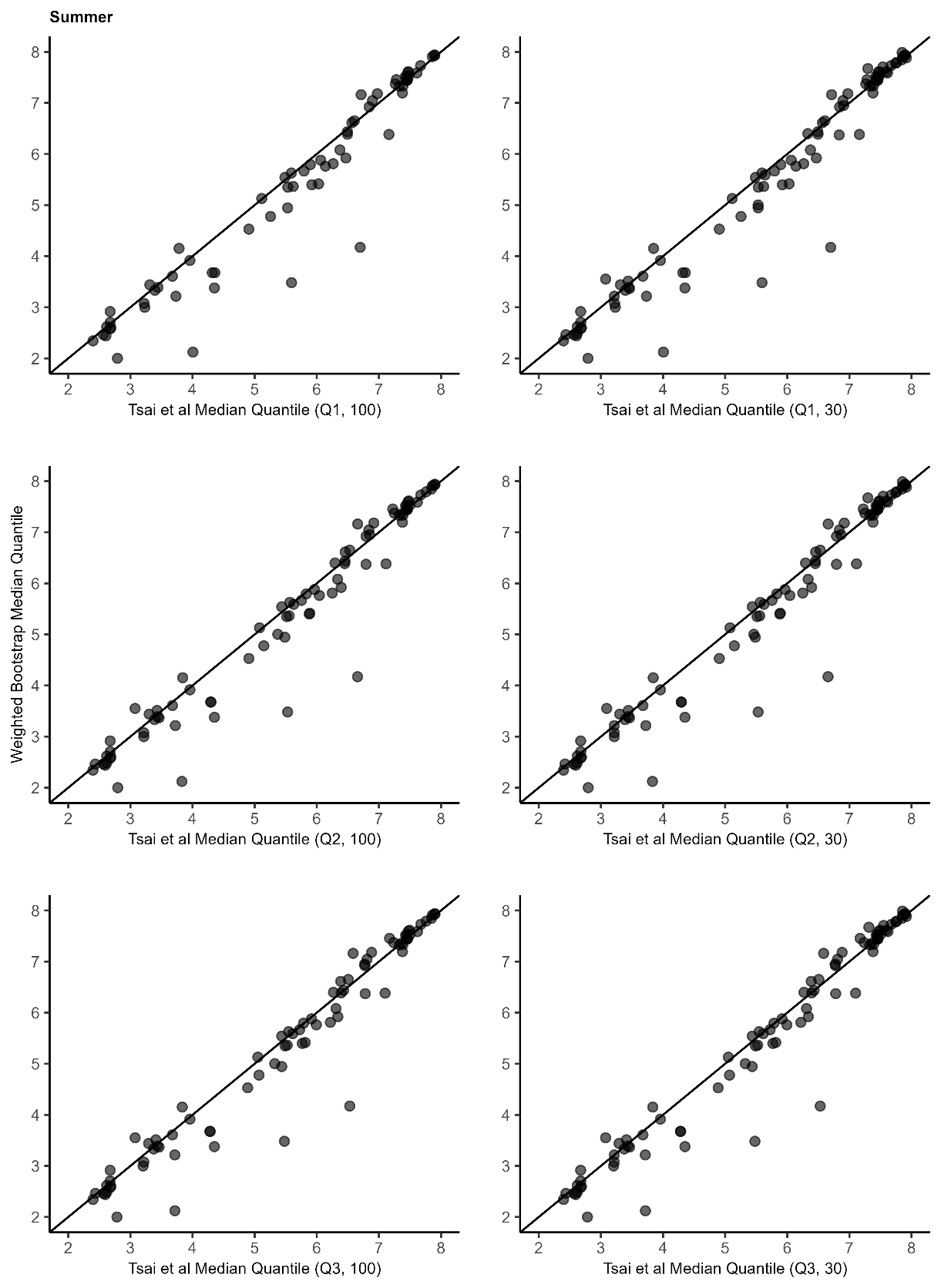

Figure S1.3: Scatter plot to compare the winter median elevation estimates of the resampling method employed in this study (Weighted Bootstrap) to the method employed by Tsai et al. (2021). Each panel is a comparison of our method to different levels of sampling effort and occurrence thresholds used by Tsai et al. (2021). Q1, Q2, Q3 denotes the first, second and third quantile of sampling effort, while 30 and 100 are the different occurrence thresholds in their analysis. A solid black 1:1 line is included to offer easier comparison between the two methods.

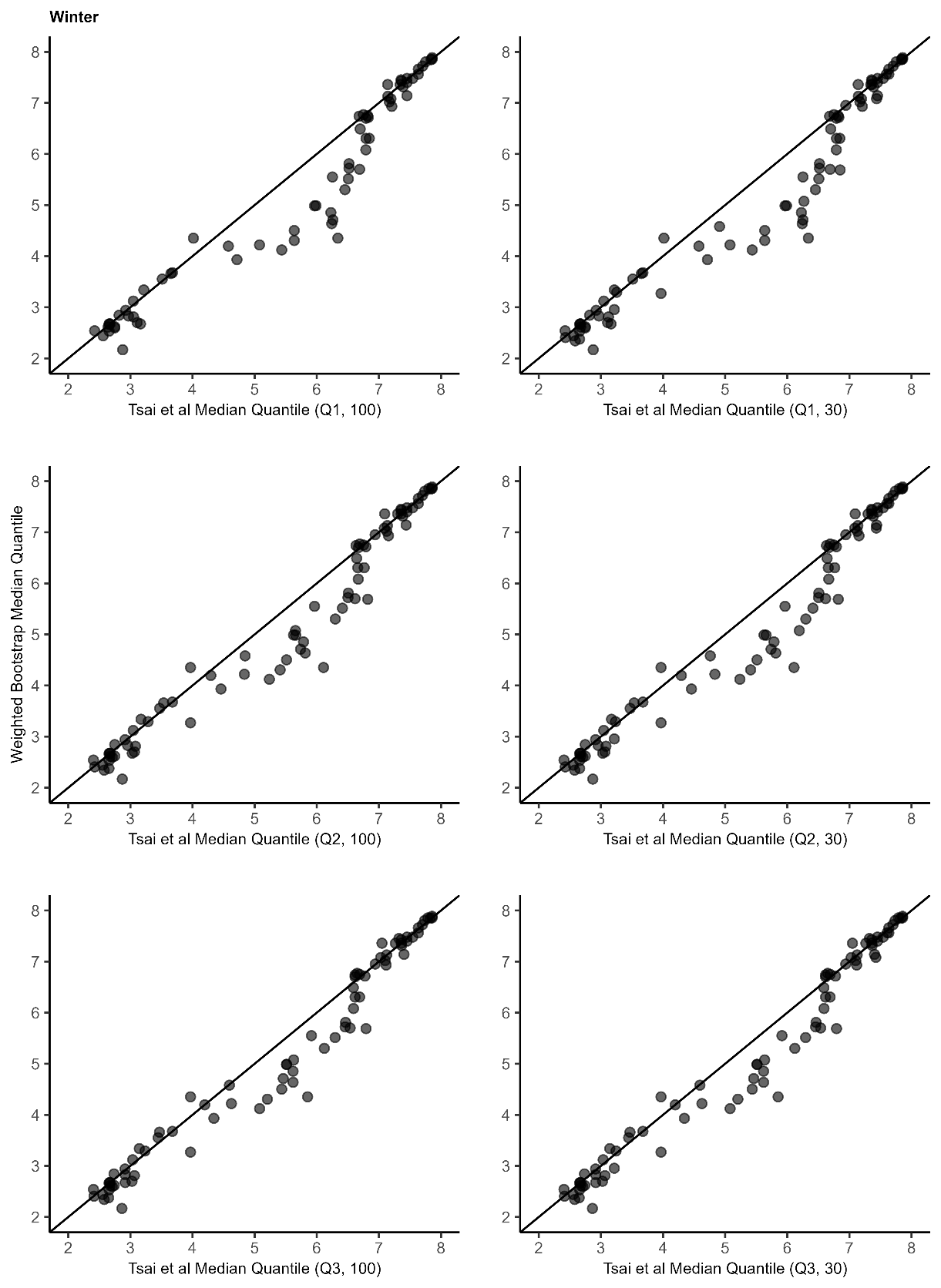

Figure S1.4: Bar plot to compare the execution time of the resampling method employed in this study (Weighted Bootstrap) versus the method employed in Tsai et al. (2021).

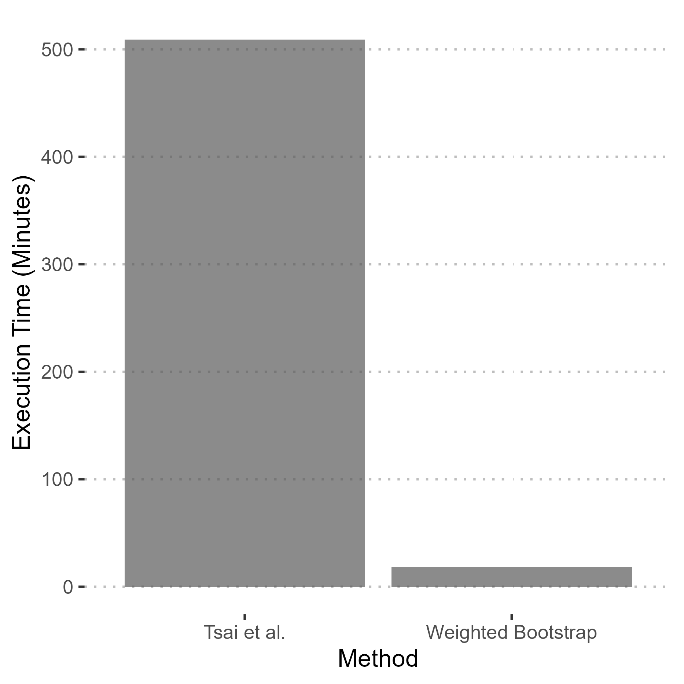

Table S1.1: Median elevation estimates for each species in this study along with their elevational migration distances for each season. A distance value in red indicates the species exhibits a significant elevational shift in that season.

| Scientific Name | Summer_median_ | Monsoon_median_ | Winter_median_ | Summer Distance | Monsoon Distance | Winter Distance |
| --- | --- | --- | --- | --- | --- | --- |
| *Columba livia* | 503.31 | 492.27 | 527.54 | -24.23 | -11.04 | 35.27 |
| *Columba elphinstonii* | 1576.21 | 1701.335 | 1526.97 | 49.24 | 125.125 | -174.365 |
| *Streptopelia decaocto* | 406.62 | 424.62 | 424.62 | -18 | 18 | 0 |
| *Spilopelia chinensis* | 927.99 | 848.97 | 776.88 | 151.11 | -79.02 | -72.09 |
| *Spilopelia senegalensis* | 640.59 | 635.86 | 619 | 21.59 | -4.73 | -16.86 |
| *Chalcophaps indica* | 658.69 | 494.79 | 689.175 | -30.485 | -163.9 | 194.385 |
| *Treron affinis* | 670.88 | 640.59 | 609.32 | 61.56 | -30.29 | -31.27 |
| *Treron phoenicopterus* | 640.59 | 703.65 | 640.59 | 0 | 63.06 | -63.06 |
| *Ducula aenea* | 439.28 | 460.12 | 94.84 | 344.44 | 20.84 | -365.28 |
| *Centropus sinensis* | 641.01 | 589.17 | 605.14 | 35.87 | -51.84 | 15.97 |
| *Centropus bengalensis* | 933.69 | 641.62 | 929.73 | 3.96 | -292.07 | 288.11 |
| *Taccocua leschenaultii* | 643.44 | 617.25 | 617.25 | 26.19 | -26.19 | 0 |
| *Phaenicophaeus viridirostris* | 576.09 | 471.56 | 549.2 | 26.89 | -104.53 | 77.64 |
| *Eudynamys scolopaceus* | 428.9 | 433.25 | 406.62 | 22.28 | 4.35 | -26.63 |
| *Cacomantis sonneratii* | 306 | 116.62 | 133.71 | 172.29 | -189.38 | 17.09 |
| *Cacomantis passerinus* | 681.11 | 643.11 | 631.95 | 49.16 | -38 | -11.16 |
| *Surniculus dicruroides* | 589.17 | 383.875 | 169.12 | 420.05 | -205.295 | -214.755 |
| *Hierococcyx varius* | 471.56 | 433.74 | 484.955 | -13.395 | -37.82 | 51.215 |
| *Cuculus micropterus* | 463.76 | 592.74 | 112.76 | 351 | 128.98 | -479.98 |
| *Harpactes fasciatus* | 542.365 | 447.325 | 495.015 | 47.35 | -95.04 | 47.69 |
| *Ocyceros birostris* | 587.53 | 586.82 | 521.81 | 65.72 | -0.71 | -65.01 |
| *Ocyceros griseus* | 745.19 | 736.18 | 644.55 | 100.64 | -9.01 | -91.63 |
| *Anthracoceros coronatus* | 180.21 | 122.2 | 129.88 | 50.33 | -58.01 | 7.68 |
| *Nyctyornis athertoni* | 874.235 | 606.38 | 640.59 | 233.645 | -267.855 | 34.21 |
| *Merops orientalis* | 580.18 | 539.425 | 564.03 | 16.15 | -40.755 | 24.605 |
| *Merops leschenaulti* | 1008.14 | 600 | 822.77 | 185.37 | -408.14 | 222.77 |
| *Coracias benghalensis* | 437.21 | 413.94 | 420.78 | 16.43 | -23.27 | 6.84 |
| *Psilopogon malabaricus* | 895.25 | 870.56 | 861.62 | 33.63 | -24.69 | -8.94 |
| *Psilopogon haemacephalus* | 640.59 | 609.37 | 587.48 | 53.11 | -31.22 | -21.89 |
| *Psilopogon zeylanicus* | 463.76 | 463.76 | 399.905 | 63.855 | 0 | -63.855 |
| *Psilopogon viridis* | 860.6 | 786.09 | 808.29 | 52.31 | -74.51 | 22.2 |
| *Picumnus innominatus* | 868.2 | 882.56 | 931.06 | -62.86 | 14.36 | 48.5 |
| *Hemicircus canente* | 464.12 | 365.29 | 491.84 | -27.72 | -98.83 | 126.55 |
| *Yungipicus nanus* | 749.32 | 594.042 | 640.59 | 108.73 | -155.278 | 46.548 |
| *Leiopicus mahrattensis* | 718.64 | 636.24 | 686.555 | 32.085 | -82.4 | 50.315 |
| *Chrysocolaptes guttacristatus* | 801.83 | 740.31 | 762.66 | 39.17 | -61.52 | 22.35 |
| *Chrysocolaptes festivus* | 640.59 | 460.12 | 640.59 | 0 | -180.47 | 180.47 |
| *Micropternus brachyurus* | 640.59 | 740.31 | 655.935 | -15.345 | 99.72 | -84.375 |
| *Dinopium javanense* | 888.74 | 902.78 | 888.275 | 0.465 | 14.04 | -14.505 |
| *Dinopium benghalense* | 475.12 | 422.78 | 425.32 | 49.8 | -52.34 | 2.54 |
| *Picus chlorolophus* | 530.82 | 512.925 | 458.09 | 72.73 | -17.895 | -54.835 |
| *Picus xanthopygaeus* | 1132.06 | 1132.06 | 1105.645 | 26.415 | 0 | -26.415 |
| *Dryocopus javensis* | 727.965 | 681.97 | 816.445 | -88.48 | -45.995 | 134.475 |
| *Psittacula eupatria* | 632.94 | 612.31 | 612.31 | 20.63 | -20.63 | 0 |
| *Psittacula krameri* | 423.71 | 425.32 | 456.97 | -33.26 | 1.61 | 31.65 |
| *Psittacula cyanocephala* | 836.55 | 788.165 | 746.94 | 89.61 | -48.385 | -41.225 |
| *Psittacula columboides* | 830.34 | 832.34 | 848.97 | -18.63 | 2 | 16.63 |
| *Loriculus vernalis* | 799.64 | 799.015 | 770.51 | 29.13 | -0.625 | -28.505 |
| *Pericrocotus cinnamomeus* | 685.84 | 640.59 | 640.59 | 45.25 | -45.25 | 0 |
| *Pericrocotus flammeus* | 812.88 | 792.575 | 808.29 | 4.59 | -20.305 | 15.715 |
| *Coracina macei* | 254.5 | 194.84 | 277.94 | -23.44 | -59.66 | 83.1 |
| *Lalage melanoptera* | 586.56 | 640.59 | 549.8 | 36.76 | 54.03 | -90.79 |
| *Oriolus chinensis* | 119.12 | 188.91 | 181.74 | -62.62 | 69.79 | -7.17 |
| *Oriolus xanthornus* | 116.62 | 122.41 | 124.43 | -7.81 | 5.79 | 2.02 |
| *Artamus fuscus* | 580.18 | 504.53 | 420.78 | 159.4 | -75.65 | -83.75 |
| *Tephrodornis sylvicola* | 531.74 | 589.17 | 645.815 | -114.075 | 57.43 | 56.645 |
| *Tephrodornis pondicerianus* | 460.12 | 287.63 | 409.41 | 50.71 | -172.49 | 121.78 |
| *Hemipus picatus* | 917.14 | 893.48 | 880.15 | 36.99 | -23.66 | -13.33 |
| *Aegithina tiphia* | 641.43 | 640.59 | 640.59 | 0.84 | -0.84 | 0 |
| *Aegithina nigrolutea* | 271.09 | 258.41 | 258.41 | 12.68 | -12.68 | 0 |
| *Rhipidura albogularis* | 945.32 | 817.38 | 769.88 | 175.44 | -127.94 | -47.5 |
| *Rhipidura aureola* | 944.94 | 822.38 | 892.14 | 52.8 | -122.56 | 69.76 |
| *Dicrurus macrocercus* | 422.11 | 424.62 | 463.76 | -41.65 | 2.51 | 39.14 |
| *Dicrurus leucophaeus* | 728.42 | 681.11 | 808.29 | -79.87 | -47.31 | 127.18 |
| *Dicrurus caerulescens* | 681.11 | 617.25 | 640.59 | 40.52 | -63.86 | 23.34 |
| *Dicrurus aeneus* | 480.76 | 406.38 | 462 | 18.76 | -74.38 | 55.62 |
| *Dicrurus hottentottus* | 647.09 | 572.785 | 612.84 | 34.25 | -74.305 | 40.055 |
| *Dicrurus paradiseus* | 635.822 | 462.31 | 565.3 | 70.523 | -173.512 | 102.99 |
| *Hypothymis azurea* | 662.345 | 635.58 | 589.17 | 73.175 | -26.765 | -46.41 |
| *Terpsiphone paradisi* | 589.17 | 589.17 | 572.44 | 16.73 | 0 | -16.73 |
| *Lanius vittatus* | 703.65 | 703.65 | 617.25 | 86.4 | 0 | -86.4 |
| *Lanius schach* | 1059.18 | 978.38 | 860.6 | 198.58 | -80.8 | -117.78 |
| *Dendrocitta vagabunda* | 370.925 | 291.94 | 375.38 | -4.455 | -78.985 | 83.44 |
| *Dendrocitta leucogastra* | 859.06 | 760.86 | 885.49 | -26.43 | -98.2 | 124.63 |
| *Corvus splendens* | 413.15 | 406.62 | 417.285 | -4.135 | -6.53 | 10.665 |
| *Corvus macrorhynchos* | 818.015 | 636.45 | 765 | 53.015 | -181.565 | 128.55 |
| *Culicicapa ceylonensis* | 1746.86 | 1647.06 | 1605.16 | 141.7 | -99.8 | -41.9 |
| *Parus cinereus* | 939.25 | 818.97 | 839.84 | 99.41 | -120.28 | 20.87 |
| *Machlolophus nuchalis* | 258.41 | 258.41 | 271.205 | -12.795 | 0 | 12.795 |
| *Machlolophus aplonotus* | 902.295 | 860.6 | 866.06 | 36.235 | -41.695 | 5.46 |
| *Ammomanes phoenicura* | 703.65 | 703.65 | 703.65 | 0 | 0 | 0 |
| *Eremopterix griseus* | 501.53 | 667.44 | 619 | -117.47 | 165.91 | -48.44 |
| *Mirafra affinis* | 474 | 474 | 463.76 | 10.24 | 0 | -10.24 |
| *Alauda gulgula* | 436.32 | 528.25 | 463.76 | -27.44 | 91.93 | -64.49 |
| *Galerida malabarica* | 801.91 | 974.045 | 711.625 | 90.285 | 172.135 | -262.42 |
| *Orthotomus sutorius* | 717.505 | 611.94 | 612.31 | 105.195 | -105.565 | 0.37 |
| *Prinia hodgsonii* | 952.31 | 758.23 | 728.42 | 223.89 | -194.08 | -29.81 |
| *Prinia sylvatica* | 516.07 | 536.47 | 617.25 | -101.18 | 20.4 | 80.78 |
| *Prinia socialis* | 703.65 | 590.46 | 619.145 | 84.505 | -113.19 | 28.685 |
| *Prinia inornata* | 617.25 | 588.71 | 548.89 | 68.36 | -28.54 | -39.82 |
| *Cisticola exilis* | 1216.62 | 988.51 | 989.94 | 226.68 | -228.11 | 1.43 |
| *Acrocephalus stentoreus* | 550.12 | 526.14 | 589.41 | -39.29 | -23.98 | 63.27 |
| *Schoenicola platyurus* | 1599.565 | 1289 | 1000.27 | 599.295 | -310.565 | -288.73 |
| *Rubigula gularis* | 306.85 | 439.28 | 351.88 | -45.03 | 132.43 | -87.4 |
| *Pycnonotus cafer* | 641.43 | 640.59 | 635.21 | 6.22 | -0.84 | -5.38 |
| *Pycnonotus jocosus* | 915.28 | 848.97 | 839.75 | 75.53 | -66.31 | -9.22 |
| *Pycnonotus xantholaemus* | 681.11 | 757.545 | 776.88 | -95.77 | 76.435 | 19.335 |
| *Pycnonotus luteolus* | 640.59 | 600.355 | 564.345 | 76.245 | -40.235 | -36.01 |
| *Chrysomma sinense* | 640.59 | 640.59 | 640.59 | 0 | 0 | 0 |
| *Zosterops palpebrosus* | 1171.76 | 1134.19 | 1125.03 | 46.73 | -37.57 | -9.16 |
| *Dumetia hyperythra* | 703.65 | 681.11 | 640.59 | 63.06 | -22.54 | -40.52 |
| *Dumetia atriceps* | 766.34 | 723.2 | 807.615 | -41.275 | -43.14 | 84.415 |
| *Pomatorhinus horsfieldii* | 1090.21 | 1088.92 | 1049.03 | 41.18 | -1.29 | -39.89 |
| *Pellorneum ruficeps* | 922.91 | 840.66 | 832.65 | 90.26 | -82.25 | -8.01 |
| *Alcippe poioicephala* | 899.665 | 836.33 | 908.39 | -8.725 | -63.335 | 72.06 |
| *Montecincla cachinnans* | 2084.97 | 2198.15 | 2217.56 | -132.59 | 113.18 | 19.41 |
| *Montecincla fairbanki* | 1619.74 | 1823.7 | 1660.15 | -40.41 | 203.96 | -163.55 |
| *Montecincla meridionalis* | 1299.97 | 1236.62 | 1237.61 | 62.36 | -63.35 | 0.99 |
| *Argya malcolmi* | 483.38 | 588.71 | 592.57 | -109.19 | 105.33 | 3.86 |
| *Argya subrufa* | 1024.755 | 950.225 | 933.74 | 91.015 | -74.53 | -16.485 |
| *Argya striata* | 619.11 | 372.76 | 519.24 | 99.87 | -246.35 | 146.48 |
| *Argya affinis* | 424.62 | 417.24 | 463.76 | -39.14 | -7.38 | 46.52 |
| *Pterorhinus delesserti* | 963.815 | 1083.655 | 1015.88 | -52.065 | 119.84 | -67.775 |
| *Sitta castanea* | 952.18 | 945.32 | 944.56 | 7.62 | -6.86 | -0.76 |
| *Sitta frontalis* | 970.05 | 935.44 | 922.85 | 47.2 | -34.61 | -12.59 |
| *Gracula indica* | 836.5 | 850.47 | 848.97 | -12.47 | 13.97 | -1.5 |
| *Sturnia pagodarum* | 918.16 | 769.88 | 728.42 | 189.74 | -148.28 | -41.46 |
| *Sturnia malabarica* | 519.235 | 470.41 | 463.76 | 55.475 | -48.825 | -6.65 |
| *Sturnia blythii* | 755.62 | 746.97 | 700.26 | 55.36 | -8.65 | -46.71 |
| *Acridotheres tristis* | 425.32 | 453.225 | 463.76 | -38.44 | 27.905 | 10.535 |
| *Acridotheres fuscus* | 1090.21 | 971.46 | 956.97 | 133.24 | -118.75 | -14.49 |
| *Turdus simillimus* | 1554.085 | 1289 | 940.12 | 613.965 | -265.085 | -348.88 |
| *Copsychus fulicatus* | 633.23 | 612.31 | 624.66 | 8.57 | -20.92 | 12.35 |
| *Copsychus saularis* | 826.38 | 734.97 | 731.535 | 94.845 | -91.41 | -3.435 |
| *Copsychus malabaricus* | 590.28 | 589.17 | 589.17 | 1.11 | -1.11 | 0 |
| *Sholicola major* | 1996.57 | 1971.06 | 2203.2 | -206.63 | -25.51 | 232.14 |
| *Sholicola albiventris* | 1972.2 | 2101.26 | 1919.62 | 52.58 | 129.06 | -181.64 |
| *Cyornis pallidipes* | 761.23 | 642.19 | 589.17 | 172.06 | -119.04 | -53.02 |
| *Cyornis rubeculoides* | 678.515 | 529.49 | 427.97 | 250.545 | -149.025 | -101.52 |
| *Cyornis tickelliae* | 754.545 | 730.045 | 681.11 | 73.435 | -24.5 | -48.935 |
| *Eumyias albicaudatus* | 1814.7 | 1811.78 | 1667.69 | 147.01 | -2.92 | -144.09 |
| *Eumyias thalassinus* | 706.22 | 817.38 | 728.42 | -22.2 | 111.16 | -88.96 |
| *Myophonus horsfieldii* | 1085.57 | 1067.85 | 986.62 | 98.95 | -17.72 | -81.23 |
| *Ficedula nigrorufa* | 1817.09 | 1907.5 | 1878.613 | -61.523 | 90.41 | -28.887 |
| *Phoenicurus ochruros* | 614.105 | 629.63 | 669.94 | -55.835 | 15.525 | 40.31 |
| *Monticola solitarius* | 839.65 | 808.29 | 823.16 | 16.49 | -31.36 | 14.87 |
| *Saxicola maurus* | 555.275 | 548.89 | 597.69 | -42.415 | -6.385 | 48.8 |
| *Oenanthe fusca* | 718.64 | 718.64 | 713.405 | 5.235 | 0 | -5.235 |
| *Dicaeum agile* | 640.59 | 602.46 | 640.59 | 0 | -38.13 | 38.13 |
| *Dicaeum erythrorhynchos* | 425.69 | 414.1 | 449.485 | -23.795 | -11.59 | 35.385 |
| *Dicaeum concolor* | 926.355 | 850.94 | 891.66 | 34.695 | -75.415 | 40.72 |
| *Leptocoma zeylonica* | 475.12 | 461.82 | 463.76 | 11.36 | -13.3 | 1.94 |
| *Leptocoma minima* | 843.22 | 850.07 | 808.29 | 34.93 | 6.85 | -41.78 |
| *Cinnyris asiaticus* | 585.91 | 604.53 | 615.74 | -29.83 | 18.62 | 11.21 |
| *Cinnyris lotenius* | 761.92 | 693.54 | 681.11 | 80.81 | -68.38 | -12.43 |
| *Aethopyga vigorsii* | 254.5 | 192.795 | 390.26 | -135.76 | -61.705 | 197.465 |
| *Arachnothera longirostra* | 748.88 | 638.22 | 751.96 | -3.08 | -110.66 | 113.74 |
| *Irena puella* | 826.69 | 751.5 | 807.44 | 19.25 | -75.19 | 55.94 |
| *Chloropsis jerdoni* | 526.82 | 522.03 | 536.47 | -9.65 | -4.79 | 14.44 |
| *Chloropsis aurifrons* | 663.41 | 640.59 | 614.095 | 49.315 | -22.82 | -26.495 |
| *Ploceus philippinus* | 612.355 | 612.31 | 619 | -6.645 | -0.045 | 6.69 |
| *Euodice malabarica* | 570.94 | 625.255 | 619 | -48.06 | 54.315 | -6.255 |
| *Lonchura punctulata* | 666.715 | 667.44 | 640.59 | 26.125 | 0.725 | -26.85 |
| *Lonchura kelaarti* | 739.69 | 819.91 | 790.37 | -50.68 | 80.22 | -29.54 |
| *Lonchura striata* | 692.38 | 742.825 | 703.65 | -11.27 | 50.445 | -39.175 |
| *Lonchura malacca* | 703.65 | 632.14 | 504.525 | 199.125 | -71.51 | -127.615 |
| *Amandava amandava* | 672.41 | 689.12 | 619 | 53.41 | 16.71 | -70.12 |
| *Passer domesticus* | 945.32 | 850.56 | 907.225 | 38.095 | -94.76 | 56.665 |
| *Gymnoris xanthocollis* | 742.76 | 945.32 | 649.37 | 93.39 | 202.56 | -295.95 |
| *Motacilla cinerea* | 1066.89 | 1081.19 | 956.88 | 110.01 | 14.3 | -124.31 |
| *Motacilla maderaspatensis* | 753.88 | 739.78 | 738.27 | 15.61 | -14.1 | -1.51 |
| *Anthus rufulus* | 779.615 | 478.455 | 621.77 | 157.845 | -301.16 | 143.315 |
| *Anthus nilghiriensis* | 2073.88 | 1891.16 | 2056.09 | 17.79 | -182.72 | 164.93 |
| *Emberiza lathami* | 828.29 | 876.24 | 728.42 | 99.87 | 47.95 | -147.82 |

Figure S1.5: Patterns of elevational migration in this study for each season; (a) – summer, (b) – monsoon, (c) winter. Arrows denote direction of movement with length corresponding to the distance of movement. Species that exhibit significant elevational movement in a particular season are in red.

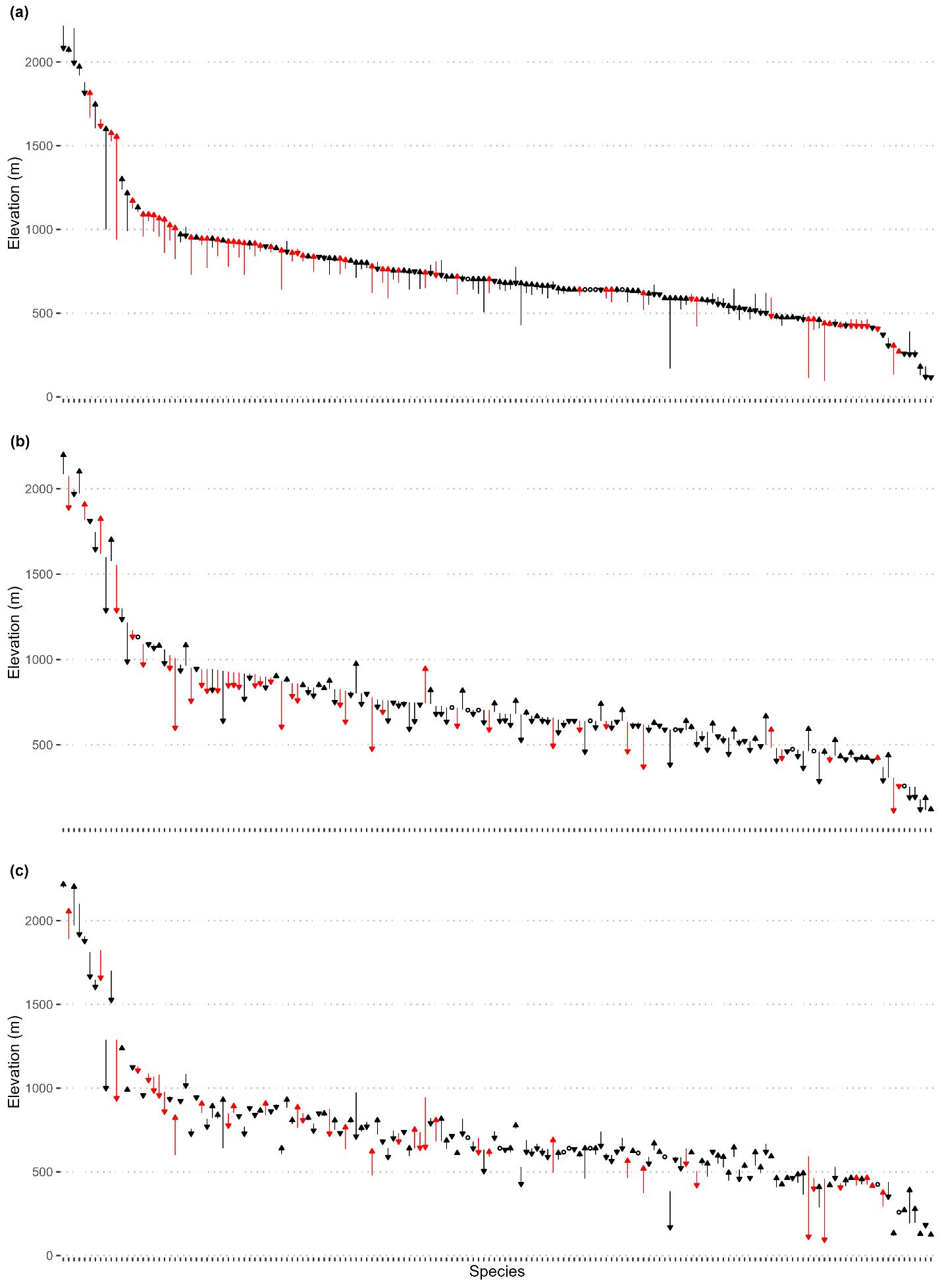

Table S1.2. The confidence set of models for each season, along with their degree of freedom (df), relative Log Likelihoods (logLik), AICc, delta AICc (*d*AICc), AIC weights (Weights), and Adjusted R^2^ (R^2^) values. An “X” present in a cell indicated that variable was not included in the model.

| (Intercept) | Diet | HWI | Mass | PTR | PMD | TTR | TMD | WTR | WMD | df | logLik | AICc | *d*AICc | Weight | R^2^ |
| --- | --- | --- | --- | --- | --- | --- | --- | --- | --- | --- | --- | --- | --- | --- | --- |
| **Summer** | | | | | | | | | | | | | | | |
| 72.175 | X | X | X | X | 15.198 | 13.956 | X | X | X | 3 | -966.416 | 1938.982 | 0 | 0.046 | 0.065 |
| 73.419 | X | X | X | X | 15.668 | 13.698 | X | 8.82 | X | 4 | -965.614 | 1939.48 | 0.497 | 0.036 | 0.074 |
| 77.8 | X | X | X | X | 19.738 | 16.329 | X | X | 11.6 | 4 | -965.639 | 1939.529 | 0.547 | 0.035 | 0.076 |
| 67.254 | X | X | 7.329 | X | 15.134 | 13.49 | X | X | X | 4 | -966.136 | 1940.524 | 1.541 | 0.021 | 0.068 |
| 67.268 | X | 7.745 | X | X | 15.491 | 13.366 | X | X | X | 4 | -966.164 | 1940.58 | 1.598 | 0.021 | 0.068 |
| 67.632 | X | X | 8.711 | X | 15.621 | 13.115 | X | 9.443 | X | 5 | -965.219 | 1940.818 | 1.836 | 0.018 | 0.078 |
| 77.574 | X | X | X | X | 11.89 | 15.169 | -7.816 | X | X | 4 | -966.284 | 1940.819 | 1.837 | 0.018 | 0.069 |
| 73.101 | X | 8.327 | X | X | 20.578 | 15.516 | X | X | 12.53 | 5 | -965.222 | 1940.823 | 1.841 | 0.018 | 0.081 |
| **Monsoon** | | | | | | | | | | | | | | | |
| 71.006 | X | X | X | 10.041 | 26.491 | 9.285 | X | X | 23.852 | 5 | -922.934 | 1856.247 | 0 | 0.058 | 0.168 |
| 71.006 | X | X | X | 8.941 | 34.376 | 11.471 | 11.526 | X | 30.794 | 6 | -921.891 | 1856.317 | 0.07 | 0.056 | 0.178 |
| 71.006 | X | X | X | X | 36.891 | 11.524 | 13.048 | X | 28.486 | 5 | -923.009 | 1856.398 | 0.151 | 0.053 | 0.167 |
| 71.006 | X | X | X | X | 28.173 | 9.016 | X | X | 20.17 | 4 | -924.346 | 1856.944 | 0.697 | 0.041 | 0.153 |
| 71.006 | X | X | X | 9.728 | 26.487 | X | X | X | 24.713 | 4 | -924.453 | 1857.157 | 0.91 | 0.037 | 0.152 |
| 71.006 | X | X | X | X | 42.26 | 13.077 | 17.477 | -8.883 | 38.508 | 6 | -922.554 | 1857.643 | 1.396 | 0.029 | 0.171 |
| 71.006 | X | X | X | X | 28.119 | X | X | X | 21.119 | 3 | -925.756 | 1857.663 | 1.416 | 0.028 | 0.138 |
| 71.006 | X | 4.668 | X | 9.911 | 27.363 | 8.896 | X | X | 25.036 | 6 | -922.567 | 1857.669 | 1.422 | 0.028 | 0.171 |
| 71.006 | X | X | X | 8.439 | 39.065 | 12.789 | 15.363 | -7.523 | 39.154 | 7 | -921.564 | 1857.846 | 1.599 | 0.026 | 0.181 |
| 71.006 | X | 3.7 | X | 8.906 | 34.58 | 11.028 | 10.813 | X | 31.304 | 7 | -921.662 | 1858.042 | 1.795 | 0.023 | 0.18 |
| 71.006 | X | 3.771 | X | X | 37.088 | 11.072 | 12.315 | X | 29.014 | 6 | -922.774 | 1858.084 | 1.836 | 0.023 | 0.169 |
| **Winter** | | | | | | | | | | | | | | | |
| 64.221 | X | 10.457 | 9.988 | 12.612 | 20.495 | X | X | X | 24.008 | 6 | -935.922 | 1884.379 | 0 | 0.047 | 0.129 |
| 64.221 | X | 9.806 | 9.963 | 12.884 | 20.375 | 7.497 | X | X | 23.148 | 7 | -935.071 | 1884.859 | 0.48 | 0.037 | 0.138 |
| 64.221 | X | 12.803 | X | 13.611 | 20.527 | X | X | X | 23.265 | 5 | -937.31 | 1885 | 0.62 | 0.035 | 0.115 |
| 64.221 | X | X | 12.379 | 12.621 | 18.638 | X | X | X | 21.751 | 5 | -937.402 | 1885.183 | 0.804 | 0.032 | 0.114 |
| 64.221 | X | X | 12.187 | 12.921 | 18.633 | 8.263 | X | X | 20.959 | 6 | -936.378 | 1885.29 | 0.911 | 0.03 | 0.125 |
| 64.221 | X | 12.144 | X | 13.881 | 20.406 | 7.529 | X | X | 22.404 | 6 | -936.466 | 1885.467 | 1.087 | 0.027 | 0.124 |
| 64.221 | X | 10.909 | 10.113 | 13.266 | 16.028 | X | -6.66 | X | 20.006 | 7 | -935.606 | 1885.929 | 1.55 | 0.022 | 0.133 |
| 64.221 | X | 10.467 | 11.064 | X | 22.636 | X | X | X | 19.535 | 5 | -937.803 | 1885.986 | 1.607 | 0.021 | 0.109 |
| 64.221 | X | 10.704 | 9.809 | 12.537 | 21.698 | X | X | -4.247 | 27.49 | 7 | -935.803 | 1886.323 | 1.944 | 0.018 | 0.131 |

*Diet = Diet, HWI = Hand-Wing Index, Mass = Body Mass, PTR = Precipitation Tolerance Range, PMD = Precipitation Median, TTR = Temperature Tolerance Range, TMD = Temperature Median, WTR = Wind-speed Tolerance Range, WMD = Wind-speed Median.
